# StrataChip: a microphysiological system capturing dynamic keratinocyte fate and mechanical transitions during human epidermal morphogenesis

**DOI:** 10.64898/2026.03.26.714483

**Authors:** Justin K. Amakor, Arvind Arul Nambi Rajan, Mageshi Kamaraj, Kyle A. Jacobs, Erica J. Hutchins, Torsten Wittmann, Matthew L. Kutys

**Author notes:** **Corresponding author:** Matthew L. Kutys.

## Abstract

Epidermal development and homeostasis require precise coordination between keratinocyte differentiation and mechanics. Still, the mechanisms integrating these processes remain poorly understood in part due to limitations of existing experimental systems. Here, we introduce StrataChip, a tractable microphysiological system that enables dynamic, multimodal interrogation of human epidermal morphogenesis. The platform integrates a media perfused dermal tissue with human epidermal keratinocytes within a microfluidic device and supports rapid epidermal stratification following establishment of an air-liquid interface. High-resolution confocal imaging and single-cell RNA-sequencing demonstrate that the StrataChip recapitulates key architectural and molecular features of human epidermis, including distinct basal, spinous, and granular layers defined by canonical differentiation markers and adhesion molecule organization. Single-cell profiling reveals transcriptionally distinct basal and spinous subpopulations, including transitional states associated with suprabasal commitment. Live 3D imaging in situ captures keratinocyte morphodynamics including basal cell delamination and asymmetric division, linking dynamic cellular behaviors to defined differentiation fates and stratification. Altogether, StrataChip provides a robust platform for a dynamic and mechanistic interrogation of how gene regulation and cell mechanics are coupled during epidermal morphogenesis.

## INTRODUCTION

The development and maintenance of tissues require coordination between cell mechanics and fate to achieve function (Collinet and Lecuit 2021). The epidermis forms a robust, impermeable barrier to external challenges, yet at the cellular scale remains dynamic to achieve homeostatic tissue maintenance (Fuchs and Raghavan 2002). To replenish the epidermis, basal keratinocytes alter their morphology and migrate towards the tissue exterior to replace cells that are shed from outer epidermal layers. During transit, keratinocyte differentiation is also carefully orchestrated with changes in keratinocyte mechanics. Breakdown of this orchestration underlies the initiation and progression of basal cell carcinoma, psoriasis, and autoimmune disorders (Amagai 2002; Simpson et al. 2011; Livshits et al. 2012; Sumigray and Lechler 2015; Broussard et al. 2021). While the importance of keratinocyte cell-cell and cell-extracellular matrix adhesions and their associated cytoskeleton to this coordination is well understood, the mechanisms by which they cooperate with gene regulation to navigate these cell state transitions are less understood. This is in part due to a lack of tractable experimental systems that allow for dynamic, multimodal interrogation of human epidermal morphogenesis (White et al. 2023).

Current investigations into epidermal morphogenesis leverage a combination of in vitro and in vivo experimental systems. Live-cell imaging techniques, such as confocal and two-photon microscopy along with genetically tractable model systems, such as *Caenorhabditis elegans*, *Drosophila melanogaster*, and vertebrate models like frog and mice, allow the assessment of candidate genes in epidermal morphogenesis within an intact organism (Schock and Perrimon 2002; Chisholm and Hsiao 2012; Box et al. 2019). Yet these non-human approaches are constrained by finite temporal imaging windows and limited mechanistic interrogation. In vitro approaches include three-dimensional (3D) human organotypic air-liquid interface cultures derived from epidermal keratinocytes and dermal fibroblasts that enable interrogation of epidermal stratification, differentiation, and barrier formation (Poumay and Coquette 2007; Malak et al. 2020; Jia and Atwood 2024). However, these models often suffer from experimental intractability, non-physiological tissue-tissue interfaces, and incompatibility with high magnification, live-cell microscopy methods.

Here, we introduce a new microphysiological model of human epidermal morphogenesis, the StrataChip, that is highly tractable, modular, and compatible with conventional live-cell confocal microscopy. Within a microfluidic device, a media perfused dermal tissue is cultured in direct contact with human epidermal keratinocytes. Introduction of an air-liquid interface stimulates the stratification of a human epidermal tissue within seven days. Microscopy and single-cell RNA-sequencing analysis reveals the StrataChip captures the identity of all epidermal cell fates, including transitional states associated with fate commitment during epidermal stratification. Furthermore, it demonstrates the spatial arrangement of adhesion and cytoskeletal machinery across stratified epidermal layers. Finally, we demonstrate the ability to capture key cellular dynamics associated with epidermal stratification by live-cell microscopy.

## RESULTS

### A microphysiological system to recapitulate epidermal morphogenesis

To study keratinocyte morphodynamics underlying epidermal morphogenesis, we engineered a microfluidic platform, termed StrataChip, in which human epidermal tissue is constructed above a fibroblast laden hydrogel approximating the dermis (Fig. 1A, B). The platform is generated by photolithography followed by poly(dimethylsiloxane) (PDMS) soft lithography and is assembled by polymerizing a type I collagen hydrogel laden with human dermal fibroblasts within a central chamber inside the PDMS mold (Polacheck et al. 2019) (Fig. 1C). The central chamber is flanked by PDMS pillars, which constrain the fibroblast-containing hydrogel and permit media diffusion from the adjacent channels. A common limitation of 3D in vitro epidermal organotypic cultures is fibroblast-driven contractility within the fibroblast-containing hydrogel. This leads to compaction of the hydrogel, requiring a period after seeding to allow the collagen hydrogel to shrink to a stable volume, which both delays and introduces technical variability during subsequent keratinocyte seeding (Bellas et al. 2012; Arnette et al. 2016). To address this technical issue, we utilized the adherent polymer polydopamine (PDA) to functionalize the surface of the PDMS chamber prior to fibroblast-containing hydrogel introduction (Park et al. 2019), followed by polymerization, and introduction of fibroblast growth media to the adjacent channels (Fig. 1D). Relative to vehicle control, which showed a 50% decrease in hydrogel volume, PDA functionalization shows no post-polymerization contraction of fibroblast containing hydrogels within the StrataChip (Fig. 1E, F). This critical step therefore promotes a stable dermal tissue mimetic that supports the subsequent culturing of human epidermal keratinocytes without delay or technical limitations associated with traditional organotypic epidermal models.

**Figure 1:**
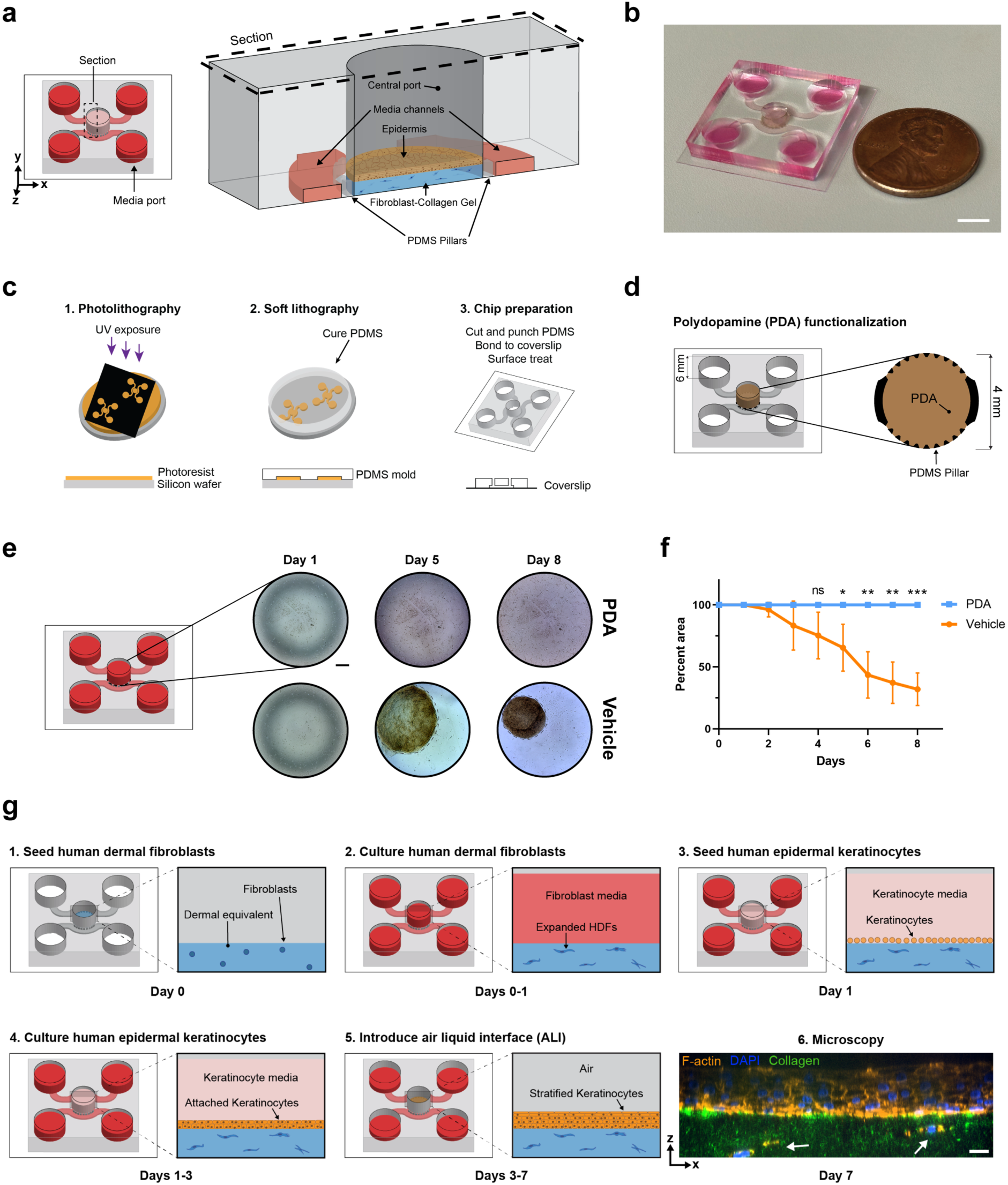
StrataChip: a microphysiological system to model epidermal morphogenesis. **(a)** 3D schematic and section rendering of the StrataChip highlighting the central port housing the epidermal tissue and dermal equivalent hydrogel surrounded by media channels. PDMS, polydimethylsiloxane. **(b)** Photograph of the StrataChip. Scale bar 5 mm. **(c)** Stepwise fabrication of StrataChip using photolithography and soft lithography to generate a PDMS device that is bonded to a coverslip **(d)** StrataChip central port treatment with polydopamine (PDA) and measurements of device features. (**e)** Time course of fibroblast-collagen hydrogel contraction, indicated by dashed lines, in the central port of PDA treated devices versus vehicle treated devices. Scale bar 0.5 mm. **(f)** Quantification of fibroblast collagen hydrogel contraction, by percent coverage of central port, in PDA treated devices versus vehicle treated devices. P = 0.01073 (Day 5); 0.00095 (Day 6); 0.00030 (Day 7); 0.00005 (Day 8) with Holm-Sidak multiple Student’s *t*- test; N = 4 independent devices for each group. **(g)** Development timeline of StrataChip: seeding of dermal equivalent (1, 2) followed by seeding of human epidermal keratinocytes (3, 4) and introduction of air-liquid interface (ALI) (5). Confocal immunofluorescence image of epidermal tissue and dermal hydrogel with fibroblasts (indicated by arrows). Visualized with phalloidin f-actin stain in orange. (6). Scale bar, 20 µm.

To generate epidermal tissue within the StrataChip, collagen-fibroblast hydrogels are cultured overnight. The next day human epidermal keratinocytes (Ker-CT) (Ramirez et al. 2003; Vaughan et al. 2009) are seeded directly on top of the underlying hydrogel in the central port and cultured for 2-3 days until confluent in the presence of keratinocyte growth media (while fibroblast growth media is compartmentalized to the lower microfluidic channels). At confluence, keratinocyte media is removed from the central port of the device to introduce an air-liquid interface (ALI), while fibroblast growth media is maintained in the microfluidic channels perfusing the dermal hydrogel (Fig. 1G). In organotypic cultures, introduction of ALI simulates key ecological features of the epidermis including the creation and maintenance of a calcium gradient and mechanical stresses that largely drive the differentiation and stratification process seen in the human epidermis (Hennings and Holbrook 1983; Malak et al. 2020). Remarkably, after just seven days in culture, the StrataChip generates a well-stratified epidermis. The generated epidermal tissue forms a physiological tissue-tissue interface with the underlying fibroblast-collagen hydrogel. Importantly, the small volume construction of the StrataChip permits detailed in situ analysis of the entire tissue via confocal microscopy (Fig. 1G).

### Single-cell transcriptomics captures key epidermal cell fates in the StrataChip

Our data suggest the StrataChip supports epidermal morphogenesis and forms a well-differentiated tissue architecture within seven days. To carefully analyze the cell dynamics and fates associated with epidermal formation and maintenance within the StrataChip, we utilized confocal microscopy and single-cell RNA-sequencing. Visualizing F-actin within developing epidermal tissues from Day 2 (prior to ALI) to Day 7 revealed progressive cell stratification culminating in epidermal tissue averaging 45 micrometers in thickness (Fig. 2A, B). In the epidermis, keratinocytes are organized into three principal layers, stratum basale, stratum spinosum, and stratum granulosum, composed of basal (BC), spinous (SC), and granular cells (GC), respectively. Each layer is defined by characteristic molecular markers and architectural features reflective of distinct functional cell fates (Koehler et al. 2011) (Fig. 2C). Keratin intermediate filaments serve as canonical markers of epidermal keratinocyte differentiation.

**Figure 2:**
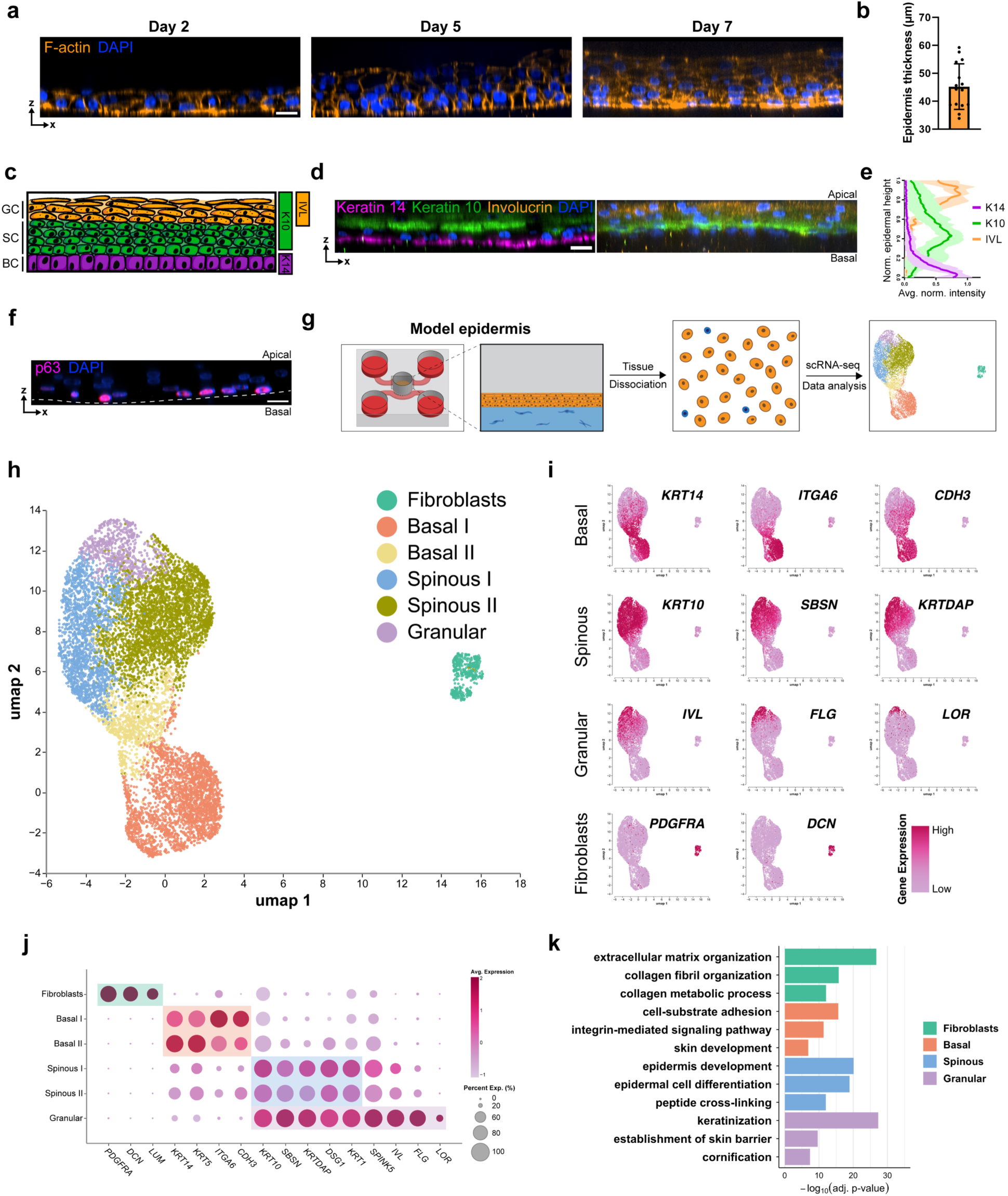
Microscopy and single-cell transcriptomics capture key epidermal cell fates in the StrataChip. **(a)** Fluorescence micrographs showing the epidermis developmental timeline in orthogonal slices at day 2, day 5 and day 7. Visualized with phalloidin f-actin stain in orange. **(b)** Bar graph of the average epidermal thickness. N = 15 independent devices. **(c)** Representative illustration of epidermis with colors representing expression of differentiation markers in basal cells (BC), spinous cells (SC) and granular cells (GC) within epidermal layers: Magenta-Keratin 14 (K14), Green-Keratin 10 (K10), Orange-Involucrin (IVL). **(d)** Orthogonal slice fluorescence micrograph of StrataChip epidermis at day 7 immunostained with differentiation markers: KRT14 (left), KRT10 (left and right), and IVL (right), as well as DNA stain DAPI. **(e)** Quantification of the layer-specific expression for epidermal differentiation markers KRT14, KRT10, and IVL. N ≥ 4 for each marker. **(f)** Orthogonal slice fluorescence micrograph of epidermis immunostained for p63. Dashed line represents cell-ECM boundary. **(g)** Flowchart of epidermal tissue dissociation, single-cell RNA-sequencing, and analysis. (**h)** Uniform manifold approximation and projection (UMAP) plot of cell types identified in StrataChip tissues. **(i)** Expression UMAP with representative gene expression for each layer-specific cell type. Dark to light pink indicates high to low gene expression. **(j)** Dot plot showing representative gene expression for each cell type in StrataChip tissues. **(k)** Three representative gene ontology (GO) terms for each keratinocyte category. All scale bars, 20 µm.

Basal keratinocytes predominantly express more compliant keratin isoforms, such as keratin 14, consistent with their proliferative and migratory behavior, whereas suprabasal spinous and granular cells express stiffer isoforms, including keratin 10, that support the mechanical barrier function of stratified tissue (Simpson et al. 2011; Wallace et al. 2012). Along with involucrin, a key component of the cornified envelope enriched in granular cells (Ekanayake-Mudiyanselage et al. 1998; Sevilla et al. 2007), these markers were used to validate cell fates in the StrataChip. Immunofluorescence analysis of epidermal tissues generated within the platform revealed three distinct stratified layers, keratin 14-positive basal layer located proximal to the underlying hydrogel, followed by keratin 10 positive spinous layer, terminating in the outermost involucrin positive granular layer (Fig. 2C-E). Furthermore, immunostaining for the stemness and self-renewal regulator, p63, revealed localization specifically to the basal layer (Fig. 2F) (Pellegrini et al. 2001).

To gain a detailed understanding of epidermal keratinocyte fate fidelity in the StrataChip, we employed single-cell RNA-sequencing. For this experiment, seven epidermal tissues were enzymatically detached from the underlying hydrogel, dissociated, and analyzed by single-cell RNA-sequencing (Fig. 2G). Single-cell clustering identified three primary cell types (basal, spinous, and granular), supporting immunofluorescence findings (Fig. 2H, I). Clusters were labeled based on established differentiation makers: basal (*KRT14*+, *ITGA6*+, *CDH3*+), spinous (*KRT10*+, *SBSN*+, *KRTDAP*+), granular (*IVL*+, *FLG*+, *LOR*+), and fibroblasts (*PDGFRA*+, *DCN*+) (Reed and Iozzo, 2002) (Fig. 2I, J). Top differentially expressed genes and subsequent gene ontology (GO) analysis of the identified broader cell types (basal, spinous, granular) further revealed distinct transcriptional features and pathways for each cell type (Fig. 2K and Supplementary Fig. 1A-D). For example, GO analysis revealed cell-substrate adhesion pathways are enriched for basal keratinocytes and cornification pathways for granular keratinocytes.

Single-cell analysis also revealed functional heterogeneity within the three principal epidermal cell types. Two transcriptionally distinct basal populations (Basal I and Basal II) were identified (Fig. 2H, J). Relative to basal II, basal I was enriched for cell-substrate adhesion programs, including multiple laminin subunits and hemidesmosome components such as *COL17A1* (Natsuga et al. 2019) (Supplementary Fig. 2A). In contrast, basal II displayed elevated expression of stress-associated keratins (*KRT16*, *KRT6A/B*) together with increased transcription of suprabasal markers, including *DSG1*, *DSP*, and *KRT10* (F. Wang et al. 2018) (Supplementary Fig. 2A). These features suggest that basal II represents a transitional population poised for commitment to the suprabasal spinous lineage (Supplementary Fig. 2E). Similarly, two transcriptionally distinct spinous populations (Spinous I and Spinous II) were also identified. Relative to spinous II, spinous I was enriched for ribosome biogenesis and translational programs, including *RPS8* and *RPL34*, as well as immune-associated transcripts such as *S100A8/9*. In contrast, spinous II showed increased expression of genes linked to signal transduction and actin cytoskeletal regulation, including *DOCK1* and *MYO1D* (Supplementary Fig. 2B). Gene ontology analysis corroborated these distinctions, with spinous I associated with ribosomal biogenesis and spinous II with GTPase signaling and actin organization pathways (Supplementary Fig. 2C, D). These functionally distinct spinous fates parallel populations described in human epidermis, specifically, a subset characterized by elevated *GRHL3* expression reported in a prior study (Wiedemann et al. 2023). Altogether, the StrataChip captures key cell fates during epidermal morphogenesis.

### Preservation of key epidermal architectural features in the StrataChip

Epidermal morphogenesis requires the coordination of cell fates with cell mechanics to achieve tissue form and function, so we next examined cell architectural features within the StrataChip. In situ fluorescence labeling of epidermal tissues for F-actin revealed distinctive cell architectures through the stratified layers (Fig. 3A). We observe the small and compact cell morphology that is characteristic of basal keratinocytes, the squamous cell morphology that is characteristic of the spinous layer of the epidermis, the flattened shape of the granular layer, and the flattened cell shape of the early corneum lacking nuclei (Fig. 3A; Supplementary Movie 1). In addition, we observe F-actin intensity heterogeneity and asymmetry across layers that has been recently described within whole mount mouse and human epidermis (Rübsam et al. 2017). Specifically, we identify the reduced F-actin intensity in the spinous layer when compared with the granular layer, with an average fold enrichment above 1.2, as well as individual cells with high F-actin intensity relative to neighboring cells (Supplementary Fig. 3A-C).

**Figure 3.**
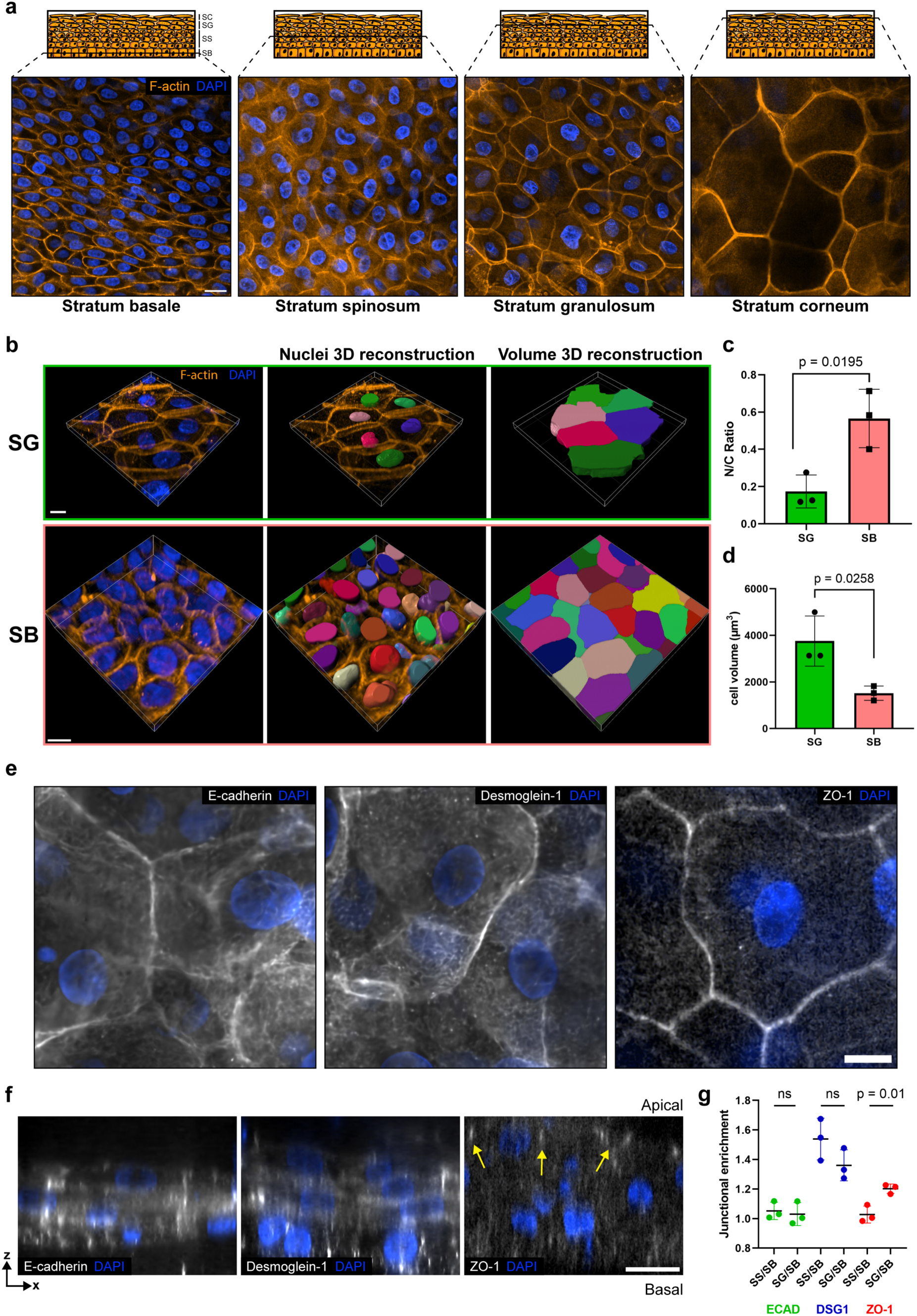
Preservation of key epidermal architectural features in the StrataChip. **(a)** Representative fluorescence images taken from the different layers of the epidermis within the StrataChip: stratum basale (SB), stratum spinosum (SS), stratum granulosum (SG), and stratum corneum (SC). Visualized with phalloidin f-actin stain in orange. Scale bar, 20 µm. **(b)** Imaris segmentation and volume analysis of epidermal sections of the SG and SB layers of the model epidermis. Scale bar, 10 µm. **(c)** N/C ratio and **(d)** cell volume quantification of Imaris cell segmentation. N = 3 independent devices, Student’s *t*- test. **(e)** Immunofluorescence micrographs of cell-cell adhesion molecules desmoglein 1, E-cadherin, and ZO-1 in granular layer of epidermis. Scale bar, 10 µm. **(f)** Orthogonal micrographs of full epidermis with immunofluorescence staining of desmoglein 1, E-cadherin, and ZO-1. Scale bar, 20 µm **(g)** Quantification of relative enrichment of cell-cell adhesion machinery within each differentiated layer normalized to the basal layer. N = 3 independent devices for each adhesion molecule, Student’s *t*- test.

Given the high-resolution 3D imaging enabled by the StrataChip, we sought to quantify the distinct architectural features of epidermal cell types using individual cell segmentation and quantification of cell volume and nuclear to cytoplasmic volumetric ratio (N/C) (Fig. 3B). Consistent with observations in vivo (Liao et al. 2013), we observe a larger N/C ratio within basal keratinocytes compared to granular keratinocytes, as well as larger cell volume of cells within the granular layer of the epidermis (Fig. 3C, D). Epidermal tissue in the StrataChip also exhibits layer-appropriate expression and spatial organization of key adhesion molecules, consistent with the defined differentiation trajectory (Fig. 3E). E-cadherin was ubiquitously expressed across all layers, reflecting its fundamental role in maintaining adherens junction integrity throughout basal, transitional, and suprabasal fates (Simpson et al. 2011; Moreci and Lechler 2020). In contrast, desmoglein-1 expression was enriched in the spinous and granular compartments relative to the basal layer, aligning with the emergence of suprabasal identity and progressive desmosome maturation during commitment (Mahoney et al. 2006). Finally, ZO-1 expression was restricted to the most apical cells consistent with tight junction assembly in terminally differentiated granular keratinocytes (Niessen 2007) (Fig. 3F, G). Together, these adhesion patterns mirror the transcriptional stratification identified by single-cell analysis and reinforce the coordinated coupling between epidermal differentiation fate, and cell and junctional architecture in the StrataChip model.

### Live 3D imaging of keratinocyte dynamics during epidermal morphogenesis

Human epidermal organotypic in vitro models are widely used for examining tissue function and fixed time point architecture, but their utility for live imaging of underlying cellular processes is still emerging (Frankart et al. 2012; Ipponjima et al. 2020). All stratifying keratinocytes originate at the lowest layer of the epidermis, the stratum basale, through multiple coordinated division and cell migration steps such as asymmetric division and delamination. Here, we investigated whether these critical processes are captured using live-cell imaging within the StrataChip. We utilized cell-permeable, fluorescent live-cell probes, FastAct and Spy-DNA, to label and visualize keratinocytes (via F-actin) and nuclear dynamics within formed epidermal tissues. This approach enabled us to visualize the distinct keratinocyte morphologies across each stratified layer and, remarkably, also revealed layer-specific cell morphodynamics within the live 3D tissue context (Supplementary Movie 2).

Focusing on early stratification stages, using 3D live imaging and image reconstruction we observed distinctive transitions of keratinocytes localized at the basal extracellular matrix interface to suprabasal layers. In one instance, we observe delamination of basal keratinocytes and subsequent intercalation into suprabasal layers (Fig. 4A; Supplementary Movie 3-5). During this process, basal keratinocytes progressively reduce cell-extracellular matrix attachment and intercalate into the upper layer independent of mitotic division. In other instances, we observe asymmetric mitotic divisions (Fig. 4B; Supplementary Movie 6, 7). Here, basal keratinocytes round and divide, with one cell maintaining basal extracellular matrix contact and the other oriented toward the suprabasal layer. In addition to early processes, we also observe nuclear degradation and DNA leakage characteristic of keratinocytes during the cornification and terminal differentiation process (Ipponjima et al. 2020). Consistent with outer cornified keratinocytes lacking nuclei in fixed tissues (Fig. 3A), live tissue imaging reveals nuclear degradation and nuclear leakage of specifically apically located nuclei within the outermost tissue layers (Fig. 4C, D; Supplementary Movie 8). Together, these observations show that live imaging in the StrataChip captures key keratinocyte morphodynamic behaviors associated with distinct differentiation stages of epidermal stratification.

**Figure 4:**
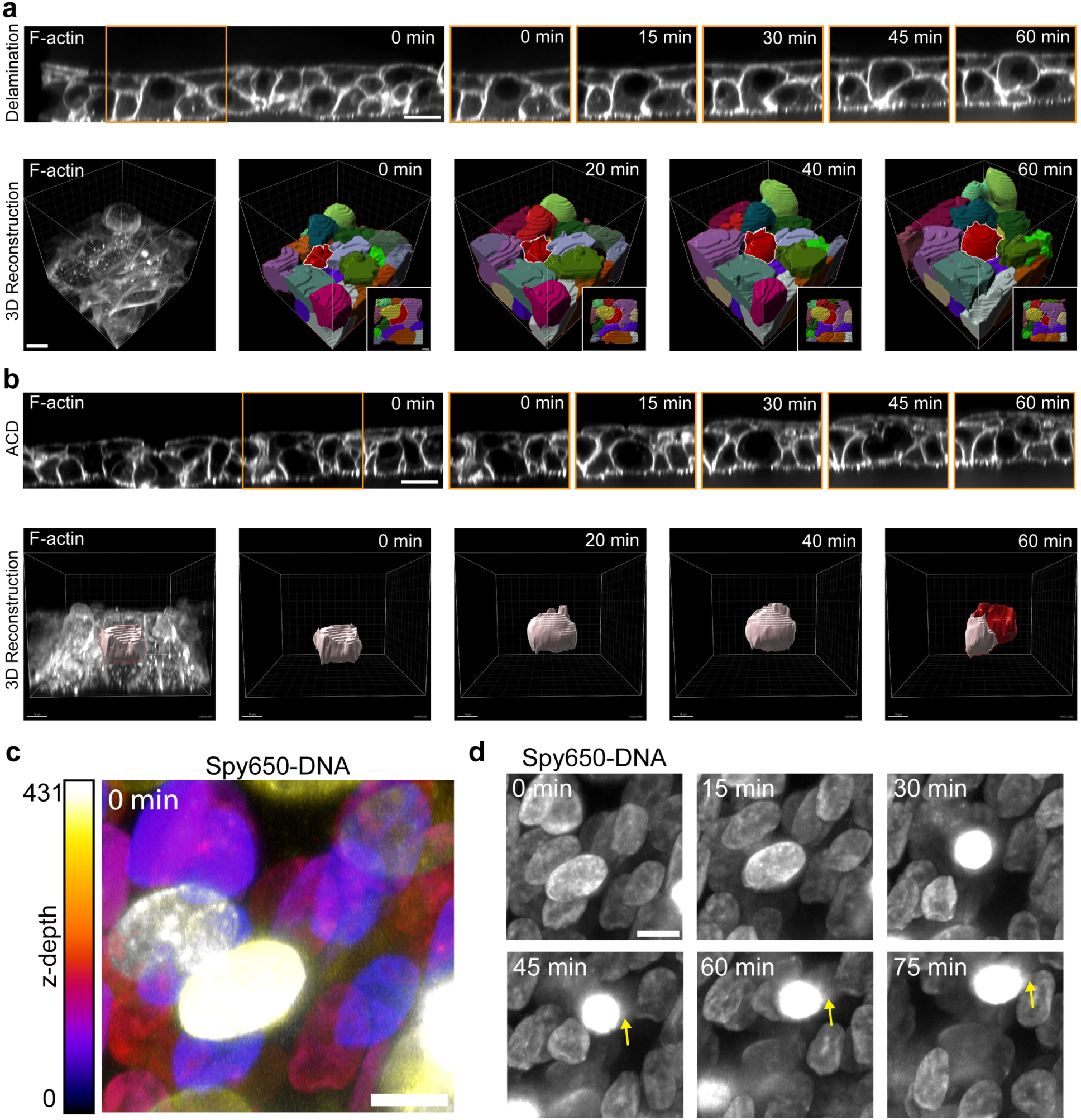
Live 3D imaging of keratinocyte dynamics during epidermal morphogenesis. **(a)** Orthogonal micrographs of FastAct-labeled live imaging of StrataChip epidermis highlighting a representative example of delamination in the developing basal layer with Imaris reconstruction and bottom point of view (POV). Red cell with white highlights is the delaminating cell in Imaris reconstruction. **(b)** Orthogonal micrographs of FastAct-labeled live imaging of StrataChip epidermis highlighting a representative asymmetric cell division (ACD) event in the basal layer with Imaris reconstruction. Red cell is the suprabasal transitioning daughter cell. **(c)** Maximum intensity projection of DAPI labeled nuclei with psuedocoloring coding for z-depth. White/yellow represents apically located nuclei and purple/black represents basally located nuclei. Scale bar, 10 µm. **(d)** Maximum intensity projection time course of DAPI labeled nuclei. Scale bar, 10 µm. Epidermal images are representative of at least 3 independent observations. Scale bar 20 µm for all orthogonal micrographs. Arrows point to DNA leakage. Scale bar 10 µm for all Imaris reconstruction.

## DISCUSSION

Organotypic human epidermal tissues grown at an air-liquid interface have served as foundational models of tissue stratification and barrier formation in vitro. Yet, these systems are often limited by technical variability, restricted optical accessibility, and reliance on bulk molecular readouts or endpoint histology, constraining analysis of keratinocyte dynamics and the cellular heterogeneity embedded within morphologically defined layers. Here, we present the StrataChip, a tractable and modular microphysiological platform that supports stable dermal-epidermal architecture, high-resolution live imaging, and single-cell transcriptomic profiling within a controlled in vitro environment. Using this system, we identify transcriptionally distinct basal and spinous subpopulations, including transitional cell states associated with fate commitment, and demonstrate the coordinated spatial arrangement of adhesion and cytoskeletal machinery across epidermal layers. Using live-cell microscopy, we further capture individual keratinocyte morphodynamics associated with stratification from the basal layer. Thus, the StrataChip extends classical organotypic models to provide a dynamic and mechanistic view of how gene regulation and cell mechanics are coupled during epidermal morphogenesis.

In addition to enabling quantitative analysis of single keratinocyte morphodynamics, the StrataChip provides a platform to interrogate how discrete keratinocyte cell states emerge, transition, and become pathologically reprogrammed. Our single-cell analysis identifies transitional basal and spinous populations enriched for stress-associated keratins (*KRT6/16*), adhesion remodeling programs, and cytoskeletal regulators, features that are frequently co-opted in epidermal inflammatory and hyperproliferative disorders such as psoriasis and early neoplastic progression (Cohen et al. 2024). The ability to combine this microphysiological system with genetic engineering, CRISPR-based perturbations and fluorescent lineage reporters, creates a tractable framework to causally link gene regulation to mechanical state transitions in real time.

For example, disease-associated mutations that perturb polarity, adhesion, or differentiation programs, such as mutations in desmosome and hemidesmosome proteins associated with Pemphigus Vulgaris and Epidermolysis Bullosa (Hoffman et al. 2025; Zhai et al. 2026), could be introduced into defined basal populations to determine whether they bias cells toward aberrant transitional states, alter delamination dynamics, or uncouple stratification from terminal differentiation. Because the platform supports live imaging alongside single-cell transcriptomics, it becomes possible to directly connect altered transcriptional trajectories with measurable changes in cell architecture, junctional organization, and migratory behavior. More broadly, this integrative approach offers a path to modeling the earliest cell state transitions that precede epidermal pathology in disorders such as basal cell carcinoma, psoriasis, and autoimmune diseases like dermatitis, enabling mechanistic dissection of how inflammatory cues, mechanical stress, and genetic lesions converge to destabilize epidermal homeostasis.

An advantage of the StrataChip is its modular architecture, which enables reconfiguration of both epithelial and stromal compartments to model a broader spectrum of stratified squamous tissues and microenvironmental complexity. Because the platform relies on a tunable hydrogel scaffold, microfluidic perfusion, and air-liquid interface control, the epithelial layer can be readily substituted with keratinocytes derived from other barrier tissues, such as oral, esophageal, or cervical epithelium, thereby extending its utility to diverse stratified squamous organs. The dermal compartment can also be readily modified to include additional cell types and matrix components. For example, endothelial cells can be incorporated to form perfusable microvascular networks, and the extracellular matrix composition tuned to better reflect tissue-specific stroma. Immune cells, including tissue-resident T cells, macrophages, or dendritic cells, can be introduced either directly into the hydrogel or through the perfused channels, enabling controlled studies of inflammatory crosstalk and immune-mediated pathology. Together, this flexibility makes the StrataChip a scalable platform for building more complex tissue environments and for defining how vascular, immune, and stromal signals influence epithelial state transitions in both homeostasis and disease.

## Data availability

Single-cell RNA-sequencing data will be made available on the NCBI Gene Expression Omnibus (GEO) or Genome Sequence Archive (GSA). All analysis code and data supporting this study are available upon request.

## Acknowledgements

The authors thank the members of the Kutys, Barber, and Wittmann labs for project input and helpful discussion. This work was supported by grants from the NIH (R35GM150987, RM1DE035338) and shared equipment grant S1OD028611-01 (Nikon CSU-W1 SoRa).

## Author contributions

J.K.A. and M.L.K. designed experiments and conceived of this study. J.K.A preformed experiments and analyzed data. M.K and K.A.J supported experimental design and performance. A.A.N.R and E.J.H supported single-cell RNA-sequencing library preparation. T.W supported light sheet imaging. J.K.A and M.L.K wrote the manuscript. All authors commented and edited the manuscript.

---

**Supplementary Figure 1:**
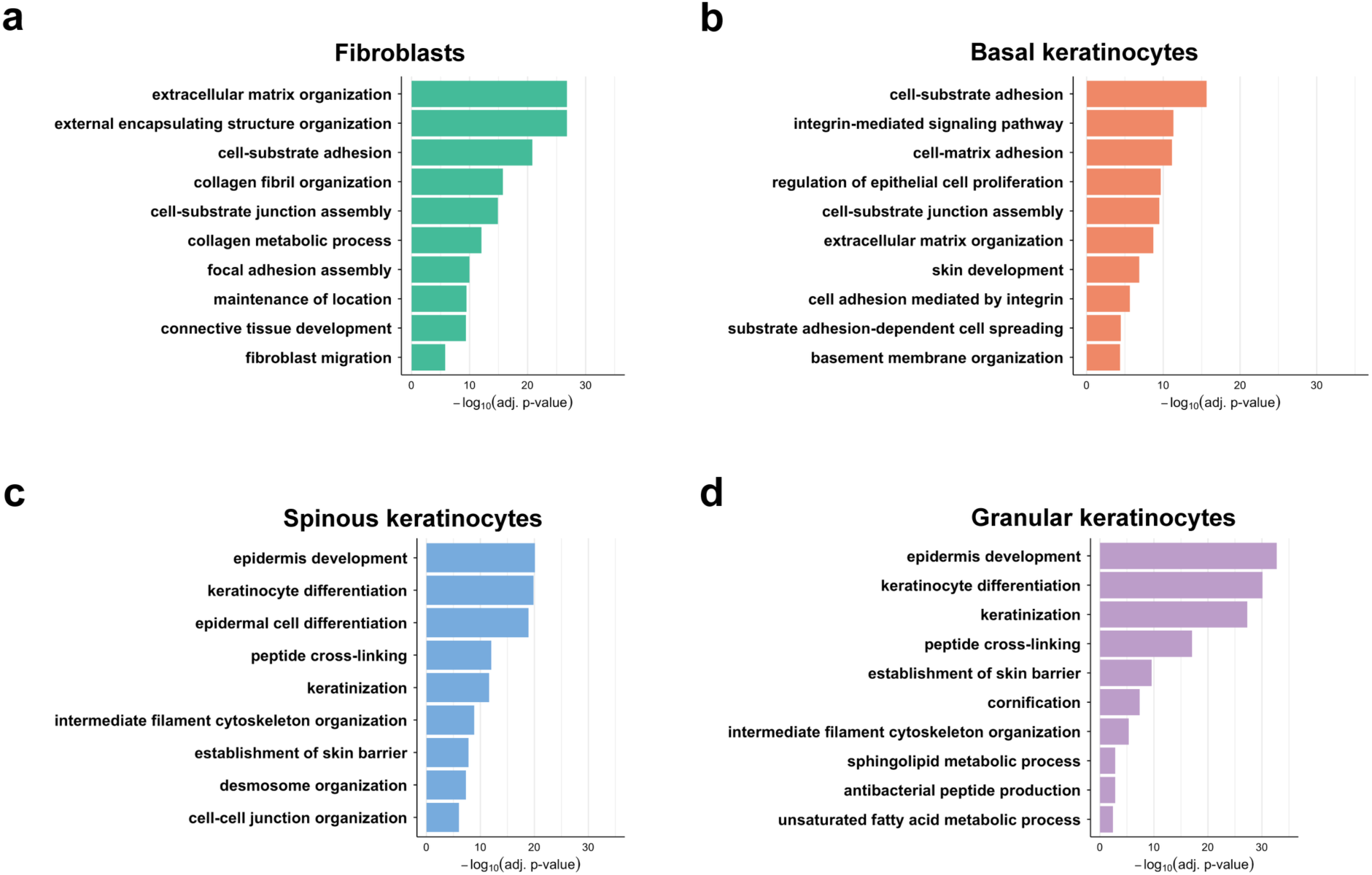
Top ten relevant GO terms from fibroblasts and each keratinocyte fate. **(a) -(d)** Top ten relevant Gene Ontology (GO) terms from the cell fates: fibroblasts and basal, spinous, and granular keratinocytes.

**Supplementary Figure 2:**
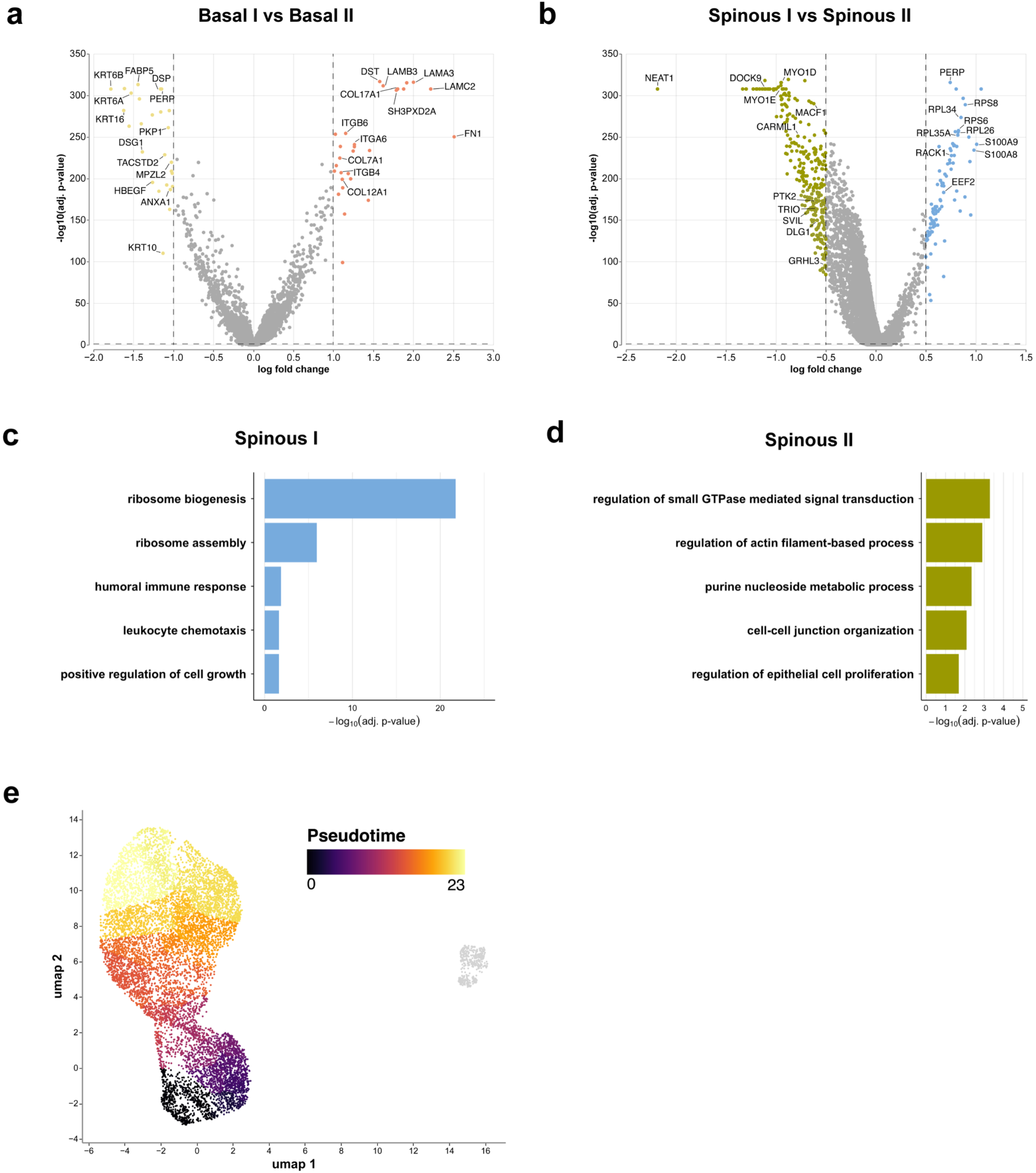
Transcriptomic differences within keratinocyte cell fates. (**a)** Differentially expressed genes between basal I versus basal II keratinocytes. **(b)** Differentially expressed genes between spinous I versus spinous II keratinocytes. **(c) and (d)** Enriched GO terms of spinous I and spinous II keratinocytes when compared with each other. **(e)** Pseudotime analysis of cell clusters from StrataChip.

**Supplementary Figure 3:**
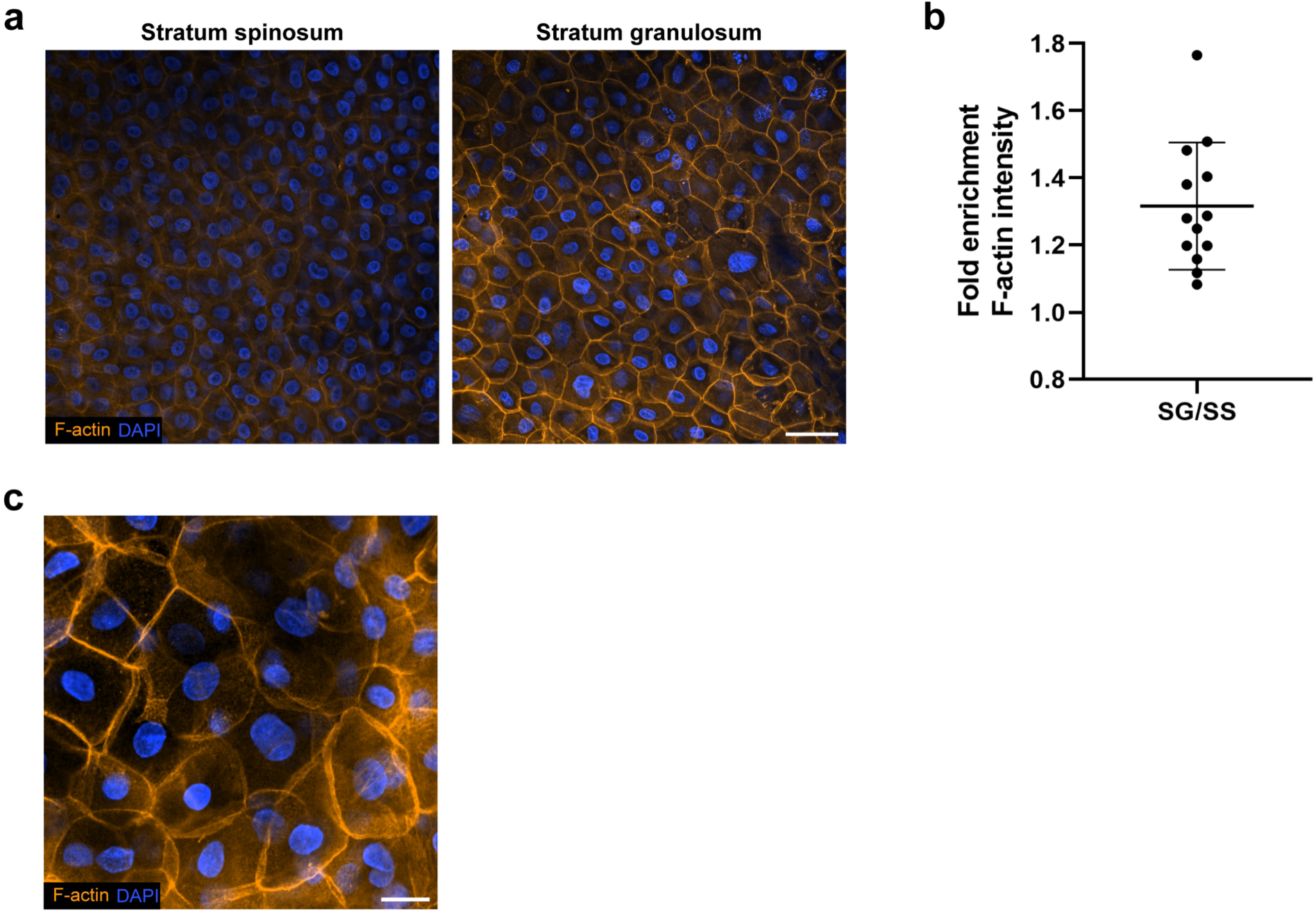
StrataChip captures keratinocyte heterogeneity and asymmetry. **(a)** Fluorescence micrographs of epidermal spinous layer and granular layer labeled for F-actin and DAPI. Scale bar 50 µm. **(b)** Normalized fold enrichment of F-actin intensity in granular layer of StrataChip epidermis. Stratum spinosum (SS) and stratum granulosum (SG). N = 13. **(c)** Fluorescence micrograph showing heterogenous F-actin intensity in granular layer of epidermis labeled with F-actin and DAPI. Scale bar 20 µm.

**Supplementary Movie 1 -** Transition through the StrataChip epidermis in z. F-actin in black and nuclei in blue. Z-step, 0.4 µm and z-depth, 49.2 µm. Displayed at 20 frames/z-steps per second. Scale bar, 50 µm.

**Supplementary Movie 2 -** Spy555-FastAct (white) and Spy650-DNA (blue) labeled keratinocytes in StrataChip visualized via time-lapse light sheet microscopy. Frames collected every 3 mins and displayed at 7 frames per second. Scale bar, 20 µm.

**Supplementary Movie 3 -** Figure 4 delamination event. Spy555-FastAct (white) labeled keratinocytes in StrataChip visualized via time-lapse confocal microscopy. Frames collected every 5 mins and displayed at 3 frames per second. Scale bar, 20 µm.

**Supplementary Movie 4 -** Figure 4 Imaris delamination event reconstruction. Displayed at 3 frames per second. Scale bar, 10 µm.

**Supplementary Movie 5 -** Figure 4 Imaris delamination event reconstruction bottom POV. displayed at 3 frames per second. Scale bar, 10 µm.

**Supplementary Movie 6 -** Figure 4 asymmetric division event. Spy555-FastAct (white) labeled keratinocytes in StrataChip visualized via time-lapse confocal microscopy. Frames collected every 5 mins and displayed at 3 frames per second. Scale bar, 20 µm.

**Supplementary Movie 7 -** Figure 4 Imaris asymmetric division event reconstruction. Displayed at 3 frames per second. Scale bar, 10 µm.

**Supplementary Movie 8 -** Maximum intensity projection of Spy650-DNA (white) labeled nuclei in StrataChip visualized via time-lapse light sheet microscopy showing nuclear degradation. Frames collected every 3 mins and displayed at 7 frames per second. Scale bar, 10 µm.

**Supplementary Table 1 –** Differentially Expressed Genes (DEGs) between cell clusters

**Supplementary Table 2 –** Gene Ontology (GO) terms for cell clusters

**Supplementary File 1 –** Adobe Illustrator mask of StrataChip

## MATERIALS AND METHODS

### Cell culture

Neonatal Human Dermal Fibroblasts (NHDFs) were obtained from Lonza. NHDFs were maintained from passages 2-7 with high glucose DMEM (Gibco) supplemented with 10% fetal bovine serum (Peak or SeraPrime), and 1% penicillin/streptomycin (Sigma) and passaged at 60-80% confluency. Human epidermal keratinocytes (Ker-CTs) cells were obtained from ATCC. Ker-CTs were maintained from passages 2-7 with KBM-Gold medium (Lonza) supplemented with KGM Gold keratinocyte growth medium BulletKit (Lonza) and passaged at 60-80% confluency. All cells were maintained in a humidified incubator at 37 °C and 5% CO2 (Thermo Fisher Scientific) and tested for mycoplasma (Applied Biological Materials). Passaged cells were counted using Trypan Blue stain (Invitrogen) and the Countess 3 automated cell counter (Invitrogen).

### Antibodies and reagents

Antibodies against Keratin 10 (RKSE60, 1:100 IF), Keratin 14 (AB_10980222, 1:100 IF), Involucrin (AB_2549923, 1:100 IF), E-Cadherin (HECD-1, 1:100 IF), ZO-1 (ZO1-1A12, 1:300 IF), Laminin (AB_2133633, 1:200 IF), Rhodamine Phalloidin (1:2,000 IF), Alexa Flour goat anti-rabbit 647 IgG (1:400 IF), Alexa Flour goat anti mouse 488 IgG (1:400 IF) were from Invitrogen. Antibodies against IL-8 (C-11, 400ng/mL WB), and DSG1 (B-11, 1:100 IF) were from Santa Cruz Biotechnology. Antibody against p63 (D9L7L, 1:400 IF) was from Cell signaling Technology. DAPI nuclear stain (1:400) was from Sigma. Spy555-FastAct (1:1,000 IF) and Spy650-DNA (1: 1,000 IF) was from Cytoskeleton Inc.

### Epidermal device Fabrication

#### Device design

Detailed device parameters and relevant dimensions are shown in figure 1D and Adobe Illustrator mask available in the supplementary files. The mask of the device was made on Adobe Illustrator. Device consists of 4 total media ports and a central port surrounded by PDMS pillars that are each 400 µm tall to contain the dermal equivalent. Each media port pair is connected by a channel that runs from one media port, alongside the central port and then to the other media port.

#### Photolithography

Silicon wafer mold was fabricated using photolithography as described in (Polacheck et al. 2019). Silicon wafer surface was cleaned with compressed air and then functionalized with a plasma cleaner (Harris Plasma) at 0.3 Torr for 5 mins at high setting. About 5 mL of SU-8 2100 (Kayaku Advance Materials) was dispensed on top of the silicon wafer and spun at 1000 rpm with a spin coater (Laurell) for a layer of about 200 µm. Wafer was soft baked on a 65 °C hot plate for 7 mins and then on a 95 °C hot plate for 90 mins. The silicon wafer was allowed to cool to room temperature for at least 15 mins. Another 5 mL of SU-8 2100 was dispensed on the wafer and spun for another 200 µm layer followed by subsequent soft baking steps at 65 °C and 95 °C and then cooled to room temperature (∼ 21 °C). Silicon wafer was patterned with ultraviolet (UV) exposure using the Alveole PRIMO system and a Nikon Ti2 microscope at 4x magnification at a power of 5 mJ/cm^2. Wafer was then post-exposure baked on a 65 °C hot plate for 5 mins then on a 95 °C hot plate for 30 mins. Silicon wafer was allowed to cool to room temperature. Unpolymerized SU-8 was removed by submerging silicon wafer in propylene glycol monomethyl ether acetate (PGMEA, Sigma) for at least 1 hr with gentle agitation. Wafer was then washed in isopropyl alcohol (-) and then dried with compressed air. Wafer was then surface treated with trichloro(1H, 1H, 2H, 2H-perfluorooctyl)silane (Sigma) by vaporization in a vacuum desiccator overnight to prevent polydimethysiloxane (PDMS Sylgard 184, Dow Corning) from binding to the wafer.

#### Soft lithography

A smooth plastic replica of the wafer was made as described in (Polacheck et al. 2019). Patterned silicon wafer was taped to the bottom of a 10 cm petri dish with double sided tape. A thoroughly mixed and degassed (with vacuum desiccator) 10:1 mixture by weight of PDMS and Sylgard 184 silicone elastomer curing agent (Dow Corning) was poured on top of the wafer. This pour was allowed to degas and was placed into a 60 °C oven overnight to polymerize. Polymerized PDMS was then cut out and used as a mold to make a plastic replica of wafer (Smooth-Cast 305, Smooth-On). Plastic wafer was used to make more copies of the device.

#### Device assembly

Individual PDMS devices were cut out from the plastic wafer with a razor blade and hole punched from the bottom with a 4 mm biopsy punch (Integra LifeSciences) for the central port and a 6 mm biopsy punch (Integra LifeSciences) for the media ports. Devices and 24 mm x 24 mm coverslips (Sigma) were then cleaned with tape to remove any dust or debris. Devices (with bottom side up) and coverslip were then placed into the plasma cleaner for 30 seconds at 0.3 Torr at high setting. Activated surfaces of devices and coverslip were then pressed together and permanently bonded by baking at 100 °C for 30 mins. Each device was typically adhered to a 6 cm dish with a small droplet of water.

#### Surface treatment

A 2 mg/mL solution of dopamine hydrochloride (Sigma) (PDA) was made with 10 mM Tris at a pH of 8.5. After initial mixture of PDA and Tris, solution was allowed to incubate for 10 mins protected from light. 20 µl of the PDA solution was then added to the central port of each device and distributed with pipet tip to ensure the PDA solution made contact with the PDMS pillars and was evenly distributed (devices were checked under microscope to make sure this was the case). The central port is then filled up with more PDA solution. Devices were incubated with PDA solution for 1 hr protected light. PDA solution was aspirated from the central port and washed (including media ports and channels) with 70% ethanol for each device. Devices were then submerged in 70% ethanol for 1 hr on a shaker. Devices were dried in a vacuum desiccator or overnight.

#### Device sterilization

Devices were sterilized with 70% ethanol washes described earlier. Devices were also sterilized right before use with UV light in a PCR chamber (Plas Labs) for 15 mins.

### Epidermal device seeding and culturing

#### Day 0: Fibroblast-collagen gel seeding

A 3 mg/mL concentration of rat tail collagen 1 (corning) was made with 10X DMEM (Thermo), 10X reconstitution buffer (RB) (1.2 g of NaHCO3 and 4.8 g of HEPES in 50 mL H_2_O), 1 N NaOH, and resuspended fibroblast solution at a pH of 7.4. DMEM and RB were mixed together on ice at a 1:10 ratio of soon to be added collagen. NaOH was then added followed by collagen to bring solution to a pH of 7.4. Collagen solution was mixed thoroughly and carefully to avoid introducing air bubbles and was then kept on ice until mixed with resuspended fibroblast solution. Fibroblasts were passaged with 1X phosphate buffer saline (PBS, Gibco) and 0.5% trypsin (Gibco) and resuspended at a desired concentration so that the final concentration of fibroblast was 6x10^4 cells/mL when added to the collagen solution. 15 µl of fibroblast-collagen solution was carefully pipetted onto the coverslip at the bottom of the central port of sterilized devices while making sure pipet tip does not make contact with the side of the central port and no air bubble was introduced. The side of the dish containing the device was gently tapped to make sure the gel was in contact with the PDMS pillars and was distributed as evenly as possible on the bottom of the central port (this was very briefly checked under the microscope). The devices were immediately placed in an incubator at 37 °C for 20 mins. NHDF media was then added to the top of the polymerized fibroblast-collagen gel and to the media ports and then drawn through the channels to connect them. PBS-soaked sterile KimWipes were then placed in each device dish to maintain humidity. Devices were then incubated at 37 °C.

For fluorescently labeled collagen, a stock solution was generated by NHS-ester conjugation of Collagen Type I, Rat Tail (Corning, 354236) to Alexa Fluor 647 (Invitrogen, A20006) as previously described (Doyle et al. 2015; Doyle 2018). The Alexa Fluor 647-labeled collagen was stored at 4°C and protected from light until use. Alexa Fluor 647-labeled collagen was added to previously calculated collagen I solution in a 19:1 volumetric ratio and mixed thoroughly.

#### Day 1: Keratinocyte seeding

NHDF media in the media ports, but not the channels, was removed and replaced with fresh media (about 250 µl per media port pair). Keratinocytes were passaged and resuspended at 2.25x10^6 cells/mL. 20 µl of media was removed from the central port and quickly replaced with 20 µl from resuspended keratinocyte-media solution. Devices were then incubated at 37 °C.

#### Day 2: Device maintenance and culturing pre-ALI

NHDF media was first removed from the media ports but not the channels. About 20 µl of Ker-CT media was removed from the central port and replaced with 30 µl of Ker-CT media (accounting for evaporation). Fresh NHDF media was then added back into the media ports (about 250 µl per media port pair). PBS-soaked sterile KimWipes were replaced with new PBS-soaked sterile KimWipes (replaced every two days).

#### Day 3: Air-Liquid Interface (ALI)

NHDF media was first removed from the media ports but not from the channels and then removed from the central port. As much media as possible was removed from the central port by tilting the devices and using a 20 µl pipet with a pre-wet (with fresh media) tip to remove any residual media. 50 µl of fresh NHDFmedia was then added into each media port pair. Devices were then incubated at 37 °C in a humidified incubator.

#### Day 4-6: Post-ALI Device maintenance and culturing

NHDF media in the media ports of the devices was replaced every 24 hrs (50 µl per media port pair). PBS-soaked sterile KimWipes were replaced when necessary.

#### Day 7: Fixation

NHDF media was removed from the media ports and the devices were fixed as described in the “Immunofluorescence” section.

### Epidermal model tissue dissociation and fixation

#### Tissue dissociation and cell isolation

Epidermal single-cell dissociation protocol was adapted from (Burja et al. 2022). Dispase digestion mix was prepared with 3 mL 1X PBS (Gibco), 0.75 mL of 9.6 U/mL dispase II (Sigma), 30 µl of 50 mM CaCl_2_, 90 µl of 1% bovine serum albumin (BSA, Fisher) in PBS, and 30 µl of 10 mg/mL DNAse I (Roche) and warmed to 37 °C. Forceps were used to gently pill the epidermis from the central port while minimizing the amount of fibroblast-collagen gel picked up. The detached epidermis was then transferred into an eppendorf tube with pre-warmed dispase digestion mix. This eppendorf tube was then shaken at 1000 rpm for 90 mins at 37 °C in a ThermoMixer F (Eppendorf). Dispase-digested epidermis was then centrifuged (Eppendorf) at 300xG at room temperature for 10 mins. The supernatant was discarded and the pellet was resuspended in a pre-warmed collagenase digestion mix. Collagenase digestion mix was prepared with 3 mL 1X PBS, Collagenase IV (Sigma), 30 µl of 100 mM CaCl_2_, 90 µl of 1% BSA, and 30 µl of 10 mg/mL DNAse I and warmed to 37 °C. The final working concentration of Collagenase IV was 1000 U/mL. The resuspended pellet was then shaken at 1000 rpm for 40 mins at 37 °C in the ThermoMixer. Collagenase-digested epidermis was then centrifuged at 300xG at room temperature for 10 mins. The supernatant was discarded and the pellet was resuspended with 500 µl of 0.25% trypsin-EDTA. The resuspended pellet was then shaken at 1000 rpm for 10 mins at 37 °C in the ThermoMixer. Trypsin-digested epidermis was then neutralized with 1000 µl of DMEM media and filtered through a 70 µm cell strainer (Falcon). Filtered cell solution was then centrifuged at 300xG at room temperature for 10 mins. The supernatant was discarded and the pellet was resuspended in 50 µl 0.2% BSA and kept on ice for Parse low input cell fixation.

#### Cell fixation

Single-cell suspension was fixed as described in the Parse Biosciences “Low Input Cell and Nuclei Fixation User Manual v1.2” and stored at -80 °C until barconding.

### Single-cell RNA-sequencing

#### Library preparation and sequencing

Single-cell libraries were prepared as described in the Parse Biosciences “Evercode WT v3 User Manual” with the Evercode Whole Transcriptome WT platform. In brief, fixed cells were captured using magnetic beads, uniquely barcoded and through split-pool combinatorial barcoding and in-cell reverse transcription, split into 8 sublibraries and then lysed. cDNA was then captured using streptavidin-coated magnetic beads. All barcoding, reverse transcription and PCR amplification reactions were performed in the Bio-Rad T100 Thermal Cycler. 6 cycles were used for cDNA amplification. Amplified cDNA was purified using KAPA Pure Beads (Roche). The concentration of purified amplified cDNA was evaluated using the Qubit dsDNA HS (High Sensitivity) Assay Kit and the size distribution was assessed using the Agilent TapeStation System using High Sensitivity D5000 ScreenTape and Reagents. Purified and amplified cDNA were fragmented, end-repaired, A-tailed, ligated to Illumina adapter sequences and then amplified again in indexing PCRs at 11 or 12 cycles based on concentration of cDNA. Fragmented cDNA were size-selected using KAPA Pure Beads. The concentration and size distribution of fragmented cDNA was checked again with the assays described above. Eight final sublibraries were sequenced on an Illumina NovaSeq 6000 at Parse Biosciences.

#### Single-cell RNA data processing and visualization

Single-cell RNA-sequencing FASTQ files were processed using Parse Biosciences computational pipeline (ParseBiosciences-Pipeline.1.3.1) and aligned to the GRCh38 reference genome. Trailmaker automatic quality control and filtering was used to remove cells with low transcripts (<1547.6), high percentage of mitochondrial reads (> 29.7%), and high doublet probability score (>0.73) from further downstream analysis. After removal of these cells, 9,765 cells remained with a median number of 8009 transcripts per cell. Data integration including log-normalization was also performed on Trailmaker using the Seurat Harmony method with the number of highly variable genes (HVG) and principle components set to 2000 and 30, respectively. Graph-based clustering was performed with the Leiden algorithm at a resolution of 0.35. Clusters were then visualized on Uniform Manifold Approximation and Projection (UMAP) embedding. Automated Leiden clusters were manually annotated based on expression profiles of known epidermal cell fate markers such as, but not limited to, KRT 14, KRT10, Involucrin, Loricrin, and previously identified epidermal cell populations (S. Wang et al. 2020; Zou et al. 2021; Polito et al. 2023; Wiedemann et al. 2023). UMAP and dot plots were generated on Trailmaker.

#### DEGs and Gene ontology analysis

Differentially expressed genes (DEGs) were identified for each cluster of interest compared to the rest of the cell sample with the presto implementation of the Wilcoxon rank sum test and auROC analysis. DEGs are listed in Supplementary Table 1. Generated DEGs were then filtered on R studio for genes with log fold change (logFC) > 0.5 (or (logFC) > 0.25 for spinous I versus spinous II GO analysis) and adjusted p-value < 0.05. Gene Ontology (GO) analysis of DEGs was done on R studio using the clusterProfiler R package and the Bioconductor homosapian genome wide annotation database (‘org.Hs.eg.db’). GO analysis was then visualized using the ggplot2 R package. Representative terms for each cluster were then selected from generated GO terms. GO terms are listed in Supplementary Table 2.

### Immunofluorescence and confocal microscopy

#### Epidermal device immunofluorescence

All incubation and washing steps in the device were done on a laboratory rocker (VWR) at room temperature unless otherwise specified. For all steps, unless otherwise stated, when all ports were filled, reagents were added to the central port first before the media ports to prevent hydrostatic pressure on the model epidermis. Media was removed from the media ports and devices were fixed by filling all ports with 4% paraformaldehyde (Electron Microscopy Sciences) prepared in 1X PBS with calcium and magnesium (0.9mM CaCl_2_ and 0.49mM MgCl_2_ or Gibco) pre-warmed to 37 °C. Devices were fixed for 35 mins. Devices (all ports) were then washed 3 times for 15 mins each with 1X PBS and then quenched with 100mM glycine (Sigma) in H_2_O for 1 hr. Devices were then washed again as described above and then permeabilized with 0.5% Triton-X (Sigma) in 1X PBS for 90 mins. Devices were then washed as described above and then blocked with 10% normal goat serum (Millipore) in 1X PBS blocking buffer for 1 hr. Primary antibodies were prepared in blocking buffer at concentrations detailed in the “Antibodies and reagents” section. Blocking buffer was removed from all ports of the device and then 5 µl of blocking buffer was added only to the central port of each device. 50 µl of diluted primary antibodies was added into each media port pair (100 µl total per device) and incubated overnight on a rocker at 4 °C. Devices (all ports) were washed 4 times for at least 15 mins each with 1X PBS. Secondary antibodies and fluorescent dyes were prepared in blocking buffer at concentrations detailed in the “Antibodies and reagents” section. Devices were incubated in secondary antibodies and fluorescent dyes as described earlier with the primary antibodies starting with the addition of 5 µl of blocking buffer to the central port and were incubated overnight on a rocker at 4 °C protected from light. Devices (all ports) were washed 4 times for at least 15 mins each with 1X PBS. Each device was filled with 1X PBS + 1% penicillin/streptomycin (Sigma) and then stored in parafilmed dishes at 4 °C until imaged.

#### Confocal microscopy

Confocal microscopy was conducted on a Yokogawa CSU-W1/SoRa spinning disk confocal with an ORCA Fusion BT sCMOS camera (Hamamatsu) controlled through NIS Elements software (Nikon) (Dema et al. 2024). An Apochromat 40x long working distance (LWD) numerical aperture (NA) 1.15 water immersion objective was use for in situ imaging of epidermal model (0.16 µm/pixel). 405 nm, 488 nm, 561 nm, and 640 nm solid-state lasers were used at powers between 25-50% (Mayo et al. 2025).

### Live-tissue microscopy

#### Confocal microscopy

Devices were incubated with NHDF and Ker-CT media with 1:2000 dilution of Spy555-FastAct at for at least 2 hr before imaging. Devices were imaged on a Yokogawa CSU-X1 spinning disk confocal custom-modified by Spectral Applied Research on a Nikon Ti-E microscope with a Evolve EMCCD camera (Photometrics) controlled by NIS Elements software (Nikon) (Stehbens et al. 2012). A Apochromat 40x long working distance (LWD) numerical aperture (NA) 1.15 water immersion objective was used with 100-mW diode-pumped solid-state (DPSS) 561 nm (Cobolt Jive) laser. Time-lapse acquisition was taken every 5 mins with a calibration of 0.09 µm/pixel. Imaging stage was enclosed within an imaging chamber (In Vivo Scientific) at 37 °C with humidified air at 5% CO_2_ (Mayo et al. 2025).

#### Light sheet microscopy

Devices were incubated with NHDF and Ker-CT media with 1:2000 dilution of Spy555-FastAct and Spy650-DNA for at least 2 hr before imaging. Devices were imaged on a Light sheet microscopy capturing ∼50 µm thick volumes was done on a custom-built oblique plane single objective OPM/SOLS microscope (Sapoznik et al. 2020) using a Nikon 40x water immersion objective as primary objective. Time-lapse acquisition was taken every 3 mins. Imaging stage was enclosed within an imaging chamber at 37 °C with humidified air at 5% CO_2_.

### Image processing and analysis

Confocal images were deconvoluted using NIS-Elements 3D deconvolution and denoised using NIS-Elements Denoise.AI software.

#### Collagen percent area contraction

Area of central port from 4x objective images was found on ImageJ along with collagen area with the polygon selection tool. Obtained values were divided to find percent area.

#### Epidermal height quantifications

Three random (x, y) coordinates were generated for each device confocal image. The thickness of the epidermis of each point was found and then averaged for each device.

#### Differentiation marker expression distribution

An orthogonal micrograph was generated for each device. The actin channel was used to draw a line segment spanning the height/thickness of the epidermis on ImageJ for each orthogonal micrograph. Intensity values of the KRT14, KRT10 and IVL channels along line segment were then generated on ImageJ and normalized to a value between 0 and 1 on R studio. All normalized intensity values were then compared by interpolation on R studio with a spacing of 0.001 from 0 to 1, where 1 is the normalized height/thickness of the epidermis measured. Interpolated values were then plotted on GraphPad Prism.

#### Imaris segmentation and quantifications

Z-stacks were converted to “.ims” files on Imaris and cropped to a ROI. Cell and nuclei segmentation was done using Imaris automatic cell and nuclei detection with relevant parameters such as nuclei and cell diameter and cortical actin thickness. Segmented cell and nuclei were then filtered by manually removing segmented cells near the edge of the ROI and segmented cells that are largely cut off by the ROI. Cell volume and N/C ratio was calculated on Imaris for remaining cells (at least 3 cells for each layer of interest).

#### Junctional enrichment analysis

The line segmentation tool (width of 10 pixels) on ImageJ was used to measure the intensity of E-cadherin, desmoglein-1, and ZO-1 along intercellular junctions across the different layers (stratum basale, stratum spinosum, and stratum granulosum) of the model epidermis for each device. At least 10 cell-cell junctions/interfaces were measured for each epidermal layer. Obtained values were then normalized to basal values as shown in Fig. 3G.

#### F-actin intensity quantification

A maximum intensity orthogonal projection was generated for each device. The line segmentation tool (width 50 pixels for granular layer and 70 pixels for spinous) was used to measure F-actin intensity in the spinous and granular layer of epidermis. Granular layer epidermal intensity was then normalized to spinous layer values to generate fold-enrichment F-actin intensity of the granular layer.

### Statistical analysis

All statistical analysis was conducted in GraphPad Prism. Data were tested for normal distribution via D’Agostino & Pearson test or Shapiro-Wilk test (alpha = 0.05) depending on sample size. Normally distributed data were analyzed via unpaired two-tailed student’s t-test. Graph error bars represent +/-standard deviation. Data analysis was not blinded, and experiments were not randomized.

