## Supplementary figures and images for "StrataChip: a microphysiological system capturing dynamic keratinocyte fate and mechanical transitions during human epidermal morphogenesis"

### Supplementary File 1

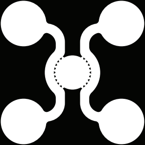
